# UGT35B1 is the principal enzyme mediating nicotine glycosylation in *Drosophila melanogaster*

**DOI:** 10.1101/2025.10.03.680272

**Authors:** Luke J. Pfannenstiel, Rachel H. Norris, Tobias Ziemke, Christophe Duplais, Nicolas Buchon, Jeffrey G. Scott

**Affiliations:** Department of Entomology, Comstock Hall, Cornell University, Ithaca, New York USA; Department of Entomology, Cornell AgriTech, Cornell University, Geneva, New York USA

**Keywords:** *Drosophila melanogaster*, UDP-glycosyltransferase, Nicotine metabolism, Detoxification, Xenobiotic glycosylation

## Abstract

Nicotine is a plant-derived pyridine alkaloid with potent neurotoxic properties. A major pathway for detoxification of nicotine in mammals is via glucuronidation to produce nicotine *N*-glucuronide, but this process in insects remains poorly understood. Using mass spectrometry, we demonstrate that *Drosophila melanogaster* detoxifies nicotine through glycosylation, producing nicotine *N*-glycoside. Given that many new agrochemicals contain pyridine rings, we also investigated the metabolism of flonicamid and imidacloprid. We detected glycosylation of flonicamid, but not imidacloprid. A targeted RNAi screen across 21 UDP-glycosyltransferases (*Ugt*s) identified *Ugt35B1* as important for survival of nicotine exposure. CRISPR-based knockout of *Ugt35B1* increases sensitivity to nicotine and flonicamid, but not to imidacloprid, nor to a structurally distinct neonicotinoid (thiamethoxam). Mass spectrometry of knockout and control flies confirms that *Ugt35B1* glycosylates nicotine, its metabolite cotinine, and flonicamid. Together these findings establish *Ugt35B1* as the principal UGT mediating nicotine detoxification in *D. melanogaster*, revealing a previously uncharacterized insect glycosylation pathway with potential implications for herbivory, insecticide detoxification and toxicology.

**Highlights:**

- *Drosophila* detoxifies nicotine by glycosylation into nicotine *N*-glycoside.
- A targeted RNAi screen identifies *Ugt35B1* as critical for nicotine survival.
- *Ugt35B1* knockout sensitizes flies to nicotine and flonicamid, but not to imidacloprid or thiamethoxam.
- First demonstration of an insect UGT mediating *in vivo* glycosylation of nicotine and cotinine.

## 1. Introduction

The interaction between insects and nicotine has long been of interest, both because of nicotine’s potent insecticidal and pharmacological activities (McIndoo, 1916; Yamamoto, 1999), as well as its occurrence in agriculturally important tobacco and other solanaceous species that serve as hosts for various insect herbivores (McKinlay et al., 1992). Beyond its natural ecological relevance, nicotine also acts by targeting nicotinic acetylcholine receptors (nAChRs), the same target site as neonicotinoid insecticides (Jeschke and Nauen, 2008). Yet, despite nicotine’s significant role in insect toxicology, the metabolic pathways by which insects detoxify this compound are not fully defined.

In mammals, nicotine metabolism proceeds primarily through three enzymatic routes: 5’-oxidation via cytochrome P450s (CYPs) to cotinine, N’-oxidation via flavin monooxygenases, and *N*-glucuronidation via UDP-glucuronosyltransferases (UGTs). All of these biotransformations increase solubility and facilitate clearance (Murphy, 2021; Yildiz, 2004). Several studies have examined the metabolism of nicotine in different insect species, commonly finding cotinine or nicotine *N*-oxide derivatives as metabolites (Wink and Theile 2002; Bass et al., 2013) (du Rand et al., 2017; Saremba et al., 2018; Self et al., 1964; Snyder et al., 1994). However, UDP-mediated conjugation has received little attention in insect metabolism, even though early studies in *Manduca sexta* reported metabolites that were “water soluble compounds which acted like conjugates” (Morris, 1983). In addition, many agrochemicals have recently been developed which contain pyridine rings (Zakharychev and Martsynkevich, 2025) suggesting that UGTs may also be of metabolic significance to these new compounds.

The UGT gene family is conserved across insects and serves dual roles: conjugating sugars to endogenous substrates involved in development and physiology (e.g., hormones, lipids), and to exogenous molecules for detoxification (Ahn et al., 2012; Kinareikina and Silivanova, 2024; Scott, 2025). Whereas mammalian UGTs predominantly utilize glucuronic acid as a sugar donor (Meech et al., 2019), insects generally use UDP-glucose to detoxify endogenous compounds including plant toxins, forming glucosides rather than glucuronides (Ahmad and Forgash, 1976; Ahn et al., 2011; Krempl et al., 2016; Robert et al., 2017; Ziemke et al., 2024) Therefore, if insects detoxify nicotine through glycosylation, the expected metabolite would be nicotine *N*-glycoside (most likely nicotine *N*-glucoside). Whether this reaction occurs *in vivo*, and which UGTs might be responsible, has remained an open question.

In this study, we investigate nicotine detoxification in *Drosophila melanogaster* with a focus on UGT-mediated metabolism. Using genetic, molecular, and analytical approaches, we define the role of glycosylation in nicotine metabolism. We show that *D. melanogaster* detoxifies nicotine through glycosylation mediated by the enzyme UGT35B1. This work addresses how insects use UGTs to defend against plant pyridine alkaloids and a structurally related insecticide, and highlights broader implications for insecticide toxicology and the evolution of xenobiotic metabolism.

## 2. Materials and Methods

### 2.1 Insects and chemicals

All *Drosophila melanogaster* stocks were maintained at 25°C under a 12:12 hour light:dark photoperiod on standard cornmeal-agar medium. The medium was prepared per liter using the following composition: 60 g cornmeal, 40 g sucrose, 50 g active dry yeast, 7 g agar, 26.5 mL Moldex (methylparaben in ethanol), and 12 mL of acid mix composed of propionic acid and phosphoric acid. Flies were reared in plastic vials or bottles with cotton plugs and maintained in incubators at constant temperature and humidity conditions.

Transgenic fly lines used in this study included TRiP-RNAi lines targeting UDP-glycosyltransferases (*Ugts*) and TRiP-CRISPR knockout (TKO) lines. The *Act5c-Cas9* strain, used for CRISPR-based gene knockout, was also employed. Unless otherwise noted, all fly stocks were obtained from the Bloomington Drosophila Stock Center (BDSC; Bloomington, IN, USA) and are listed in Supplementary Table 1.

Chemicals used in this study were: (–)-nicotine (98% purity; Sigma Aldrich, St. Louis, MO, USA), (–)-cotinine (98% purity; AA Blocks Inc., San Diego, CA, USA), flonicamid (99.5% purity; Chem Service, West Chester, PA, USA), thiamethoxam (99.5% purity; Chem Service) and imidacloprid (technical grade, Taizhou Crene Biotechnology Company, Zhejiang, China). All compounds were stored and handled according to manufacturer specifications and diluted in aqueous solutions for bioassays as described below.

### 2.2 Bioassays

Genetic crosses were performed to generate flies for use in RNA interference (RNAi) and CRISPR-based knockout bioassays. For RNAi experiments, 10 virgin females from Gal4 driver lines were crossed to five males from individual TRiP-RNAi lines targeting specific *Ugt* genes. Crosses were maintained for 48 hours at 25°C to allow mating, after which adult flies were removed. The resulting F_1_ progeny were allowed to develop, and 3-to 5-day-old F_1_ females were collected and used for toxicological assays. For CRISPR-based knockout experiments, 10 virgin females from the Act5c-Cas9 line were crossed to five males from the *Ugt35B1* TKO (TRiP-CRISPR) line. Crosses were similarly maintained for 48 hours before removing the adults. F_1_ females were collected after emergence and aged for 3 to 5 days prior to bioassay.

Toxicity assays for imidacloprid, nicotine, and thiamethoxam were performed using 20 mL glass scintillation vials (Wheaton Scientific, Millville, NJ, USA), each with an interior surface area of 38.6 cm². Test compounds were dissolved in a 10% (w/v) sucrose solution to achieve the desired concentrations. For each replicate, 20 female flies (3–5 days old) were transferred into a vial. A sterile cotton ball covered with a piece of nylon tulle was inserted into the mouth of each vial, and 100 µL of the agrochemical solution was applied evenly to the cotton ball using a syringe. The vials were laid horizontally and incubated at 25°C for 24 hours. After exposure, mortality was assessed by scoring the number of flies that were ataxic or unresponsive to gentle tapping. A minimum of three biological replicates (each from a different generational cohort) were tested for each insecticide.

Given that flonicamid did not result in mortality at any of the concentrations tested, its sublethal neurotoxic effects were instead evaluated via a negative geotaxis (climbing) assay (Qiao et al., 2022). Groups of 10 female flies (3–5 days old) were placed into 15 mL polypropylene Falcon tubes. Each tube was sealed with a cotton plug covered in nylon tulle, onto which 100 µL of flonicamid (dissolved in 10% sucrose solution) was applied. Tubes were incubated horizontally at 25°C for 24 hours. After exposure, flies were tapped to the bottom of the tube, and the number of flies that climbed above the 2 mL mark within 30 seconds was recorded as a measure of locomotor recovery. This procedure was repeated across six biological replicates (each a different generational cohort).

### 2.3 Preparation of samples for LC-MS

Female flies from the Canton-S strain or *Ugt35B1* TKO crosses were exposed to nicotine or cotinine as stated above. Twenty surviving flies were collected after 24 h of exposure and stored at −80 °C. Whole-body fly samples and frass were first freeze-dried to remove moisture and stabilize metabolites. Dried whole body samples were then suspended in 0.5 mL of methanol containing 0.1% formic acid. Homogenization was performed using a bead ruptor instrument (Omni International, Kennesaw GA, USA) equipped with stainless steel beads, ensuring thorough disruption of tissue and release of metabolites. Following homogenization, the samples were filtered using polytetrafluoroethylene (PTFE) filter vials with a 0.2 µm pore size to remove debris. Vials containing frass were rinsed with 0.5 mL of methanol containing 0.1% formic acid then filtered. These resulting filtrates were transferred into 200 µL glass insert vials compatible with autosampler racks and stored at –20°C until liquid chromatography-mass spectrometry (LC-MS) analysis.

### 2.4 Quantification of nicotine and its derivatives

Quantification of nicotine and its metabolic derivatives was conducted using reversed-phase liquid chromatography coupled to high-resolution mass spectrometry (LC-HRMS). Methanolic extracts were injected onto an Agilent InfinityLab Poroshell 120 Bonus-RP C18 column (150 mm × 3.0 mm, particle size 2.7 µm), maintained at 40°C. The mobile phase consisted of water (solvent A) and acetonitrile (solvent B) both containing 0.1% formic acid, and separation was achieved using the following gradient: 0% B at 0 minutes, held for 2 minutes; a linear increase to 95% B from 2 to 14 minutes; followed by 100% B from 14 to 17 minutes; and a final 3-minute post-run at 100% B. The flow rate was maintained at 0.6 mL/min throughout the run. Chromatographic separation was performed on an Agilent 1260 Infinity II LC system connected to an Agilent 6545 quadrupole time-of-flight (Q-TOF) mass spectrometer. Mass spectrometric detection was conducted in positive electrospray ionization (ESI) mode, scanning a mass-to-charge (m/z) range of 100–900. Instrument settings were optimized for nicotine detection and included a gas temperature of 225°C, drying gas flow of 10 L/min, nebulizer pressure of 35 psi, sheath gas temperature of 325°C, and sheath gas flow rate of 11 L/min.

Quantification was performed using Agilent MassHunter Quantitative Analysis Software. Calibration curves were generated from standard solutions of nicotine and cotinine (dissolved in methanol at concentrations of 10 ng/mL, 100 ng/mL, and 1 µg/mL. Each concentration was injected in triplicate. These calibration curves were used for the absolute quantification of nicotine and cotinine in biological samples. Glycosylated derivatives of nicotine and cotinine were quantified using a semi-quantitative approach, with concentrations estimated based on the established calibration curves for the corresponding parent compounds. Final results are expressed as nanomoles (nmol) or nanograms (ng) of compound per milligram of dry insect weight.

Structure elucidation of nicotine, cotinine, and their derivatives was performed by data-dependent analysis (DDA) tandem mass spectrometry (MS/MS) using the same LC-HRMS system and chromatographic conditions described above. MS/MS analyses were acquired in Auto MS/MS mode with a fixed collision energy of 30 eV, scanning from 40–700 m/z. Acquisition was set to a 1 s cycle time with a spectral rate of 1 Hz, and active exclusion limited fragmentation to the most intense precursor ion per cycle. All spectra and structural data can be found in SI Fig S1-S6 and Table S2.

## 3. Results

### 3.1. D. melanogaster glycosylates nicotine, cotinine and flonicamid, but not imidacloprid

To investigate how *Drosophila melanogaster* metabolizes nicotine, we exposed wild-type (Canton-S) flies to nicotine and analyzed whole-body tissues and holding vials from surviving individuals. Using HPLC-HRMS, we detected and quantified nicotine, its primary P450-mediated oxidative metabolite cotinine, and their potential glycosyl conjugates. In nicotine-exposed flies, we detected nicotine *N-*glycoside as the major metabolite in whole-body extracts, along with cotinine and cotinine *N*-glycoside, indicating that *D. melanogaster* actively glycosylates nicotine (Fig. 1A). Although our mass spectrometry data cannot distinguish between sugars with identical masses, we hypothesize glucose as a the donor, as it is the most commonly utilized sugar across seven UGTs in *Tetranycus urticae* (Snoeck et al., 2019), and in *Spodoptera* species (Wouters et al., 2014). All three metabolites, nicotine *N*-glycoside, cotinine, and cotinine *N*-glycoside, were also present in the frass (Fig. 1B), confirming that these metabolites are excreted.

**Fig. 1.**
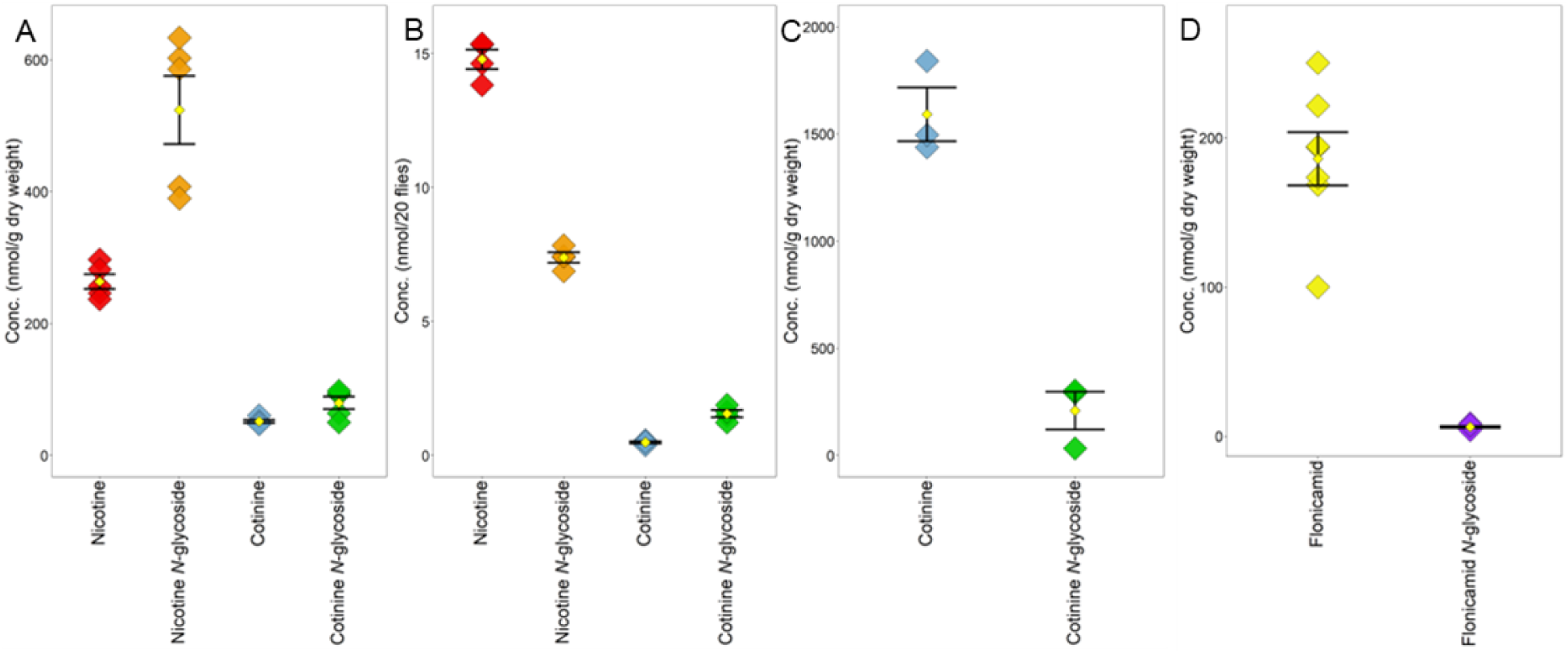
The nicotine, nicotine *N*-glucoside, cotinine and cotinine *N*-glucoside are found in nicotine exposed flies as well as in the holding vial. Dot plots showing the concentrations of nicotine and its metabolites in nicotine exposed Canton-S whole body fly samples (A) and their holding vials (B), in cotinine (C) and flonicamid (D) exposed whole body fly samples. The weight of the frass was too low to accurately measure, so its concentration is given in nmol per 20 flies.

To determine whether glycosylation occurs on other pyridine containing compounds we exposed flies to cotinine, imidacloprid and flonicamid and analyzed whole-body tissues. We detected *N*-glycoside metabolites of cotinine (Fig. 1C) and flonicamid (Fig. 1D), but not imidacloprid. Together, these results demonstrate that *D. melanogaster* UGTs can convert both nicotine and cotinine into glycosylated metabolites, supporting a role for UDP-glycosyltransferases (UGTs) in insect nicotine metabolism, potentially analogous to UGT-mediated glucuronidation in mammals. In addition, *D. melanogaster* UGTs can glycosylate flonicamid, but not imidacloprid.

### 3.2 A large-scale RNAi screen identifies *Ugt35B1* as critical for survival following nicotine exposure

In mammals, nicotine glycosylation is mediated by UDP-glycosyltransferases (UGTs), prompting us to hypothesize that one or more *D. melanogaster* UGTs could similarly catalyze the glycosylation of nicotine. As a first step toward determining the physiological relevance of this modification in insects, we set out to identify which *D. melanogaster* UGTs are involved in nicotine metabolism. To this end, we conducted a targeted RNAi screen using *Gal4/UAS*-driven knockdown of individual UGTs, followed by nicotine toxicity assays. Of the 35 *Ugt*s in the *D. melanogaster* genome, RNAi lines were available for 21 (Fig. 2A). Each of these lines was exposed to nicotine, and mortality was compared to appropriate genetic controls. The majority of RNAi knockdowns showed no significant change in nicotine sensitivity (Fig. 2B). However, knockdown of *Ugt35B1* resulted in a striking increase in nicotine-induced mortality (Fig. 2B). To validate this result additional replicates were done, and these confirmed that RNAi of *Ugt35B1* exhibited a robust increase in mortality following nicotine exposure (Fig. 2C).

**Fig. 2.**
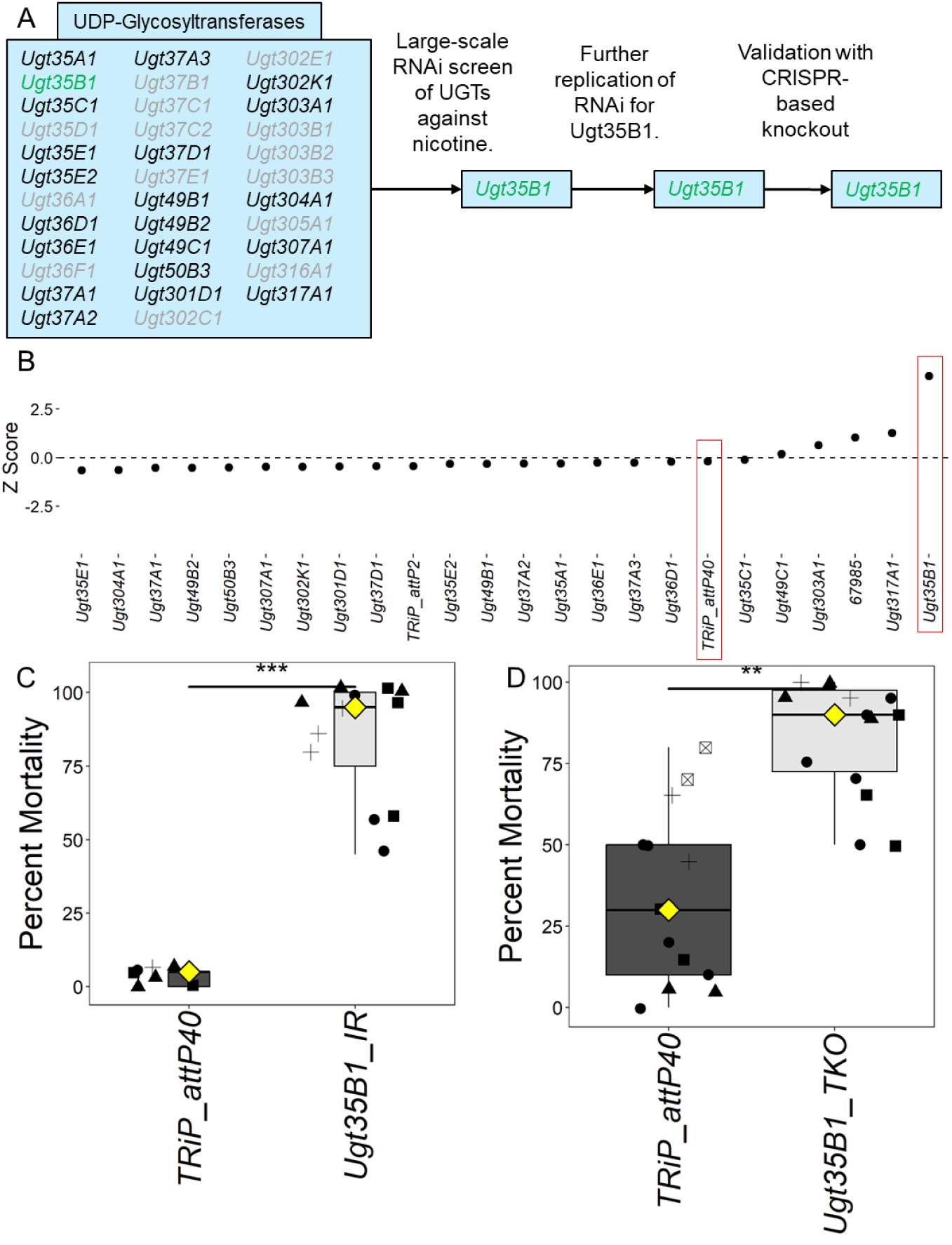
Knockdown of *Ugt35B1* significantly increases mortality after nicotine exposure. UGTs screened via RNAi for changes in response to nicotine are shown in the schematic of the screen (A). Genes in gray did not have RNAi lines available at the time of the screen, and genes in green showed an increase in nicotine toxicity. The z-score for each RNAi line and control are graphed (B). Highlighted in red are *Ugt35B1* and its control. Box plots are shown for further replication of RNAi (C) and CRISPR-based knockout (D) of *Ugt35B1* in response to nicotine. Asterisks indicate significance between knockdown/knockout and control lines. *p < 0.05, **p < 0.01, ***p < 0.001.

To confirm that *Ugt35B1* is indeed required for survival under nicotine stress, we generated a CRISPR-based whole-body knockout. These mutants also displayed significantly elevated mortality after nicotine exposure (Fig. 2D), mirroring the effect seen with the more effective RNAi line. Together, these findings identify *Ugt35B1* as a key player in *Drosophila*’s response to nicotine exposure and suggest it may directly mediate glycosylation of nicotine, contributing to detoxification and survival.

### 3.3 *Ugt35B1* is required for the glycosylation of nicotine

To directly test whether UGT35B1 catalyzes the glycosylation of nicotine and/or cotinine, we compared LC-HRMS metabolite profiles of nicotine- or cotinine-exposed background control and *Ugt35B1* knockout flies. In controls, glycosylated derivatives of both nicotine and cotinine were readily detected, consistent with our earlier observations (Fig. 3). However, in *Ugt35B1* knockout flies, levels of both nicotine *N*-glycoside and cotinine *N*-glycoside were dramatically reduced, approaching undetectable levels.

**Fig. 3.**
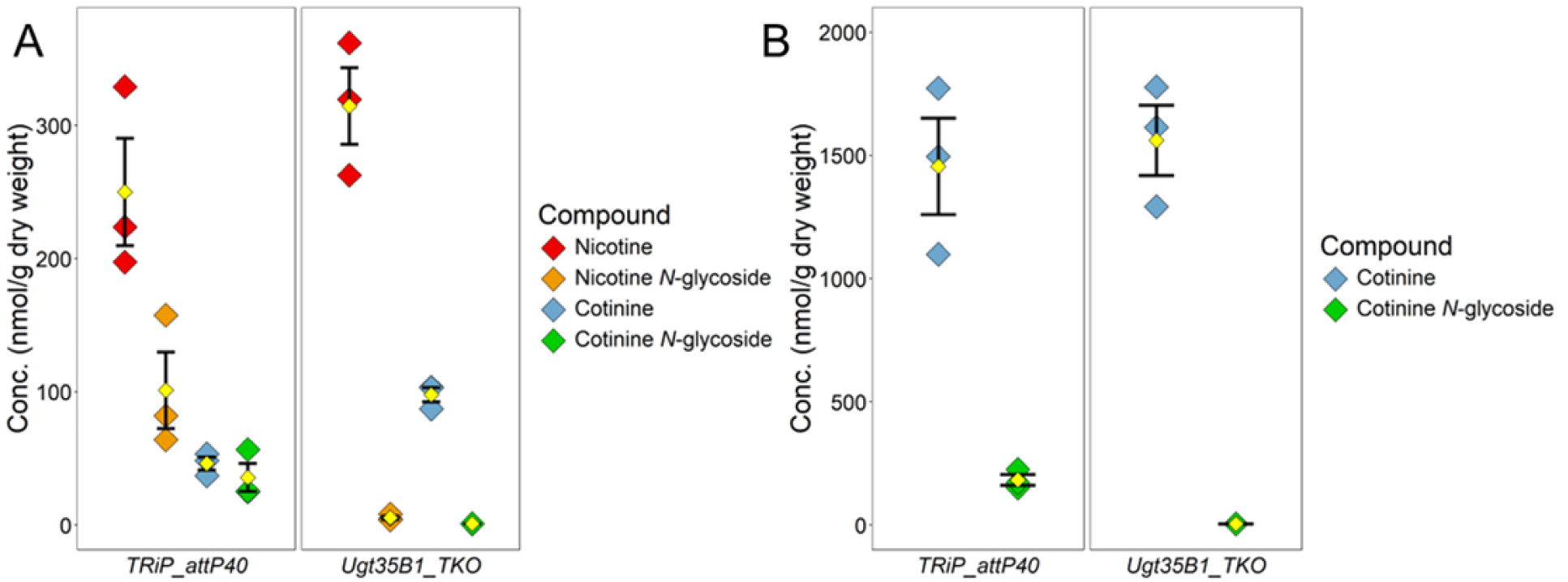
*Ugt35B1* knockout prevents nearly all nicotine and cotinine glycosylation in *Drosophila*. *Ugt35B1* TKO and control flies were exposed to nicotine (A) or cotinine (B). Dot plots show the concentrations of nicotine and its metabolites after exposure.

These findings demonstrate that *Ugt35B1* is essential for the formation of most of the glycosylated nicotine metabolites *in vivo*. The near-complete loss of these conjugates in the absence of UGT35B1, coupled with the increased mortality observed in knockout animals, establishes a causal link between glycosylation and nicotine tolerance. While trace amounts of glycosylated products were still detectable in knockout flies, suggesting that other UGTs may contribute minor activity, the magnitude of the reduction confirms that UGT35B1 is the primary enzyme responsible for this detoxification pathway.

Together, this final result unifies our physiological, genetic, and biochemical data to reveal a clear mechanistic basis for nicotine detoxification in *D. melanogaster*. By catalyzing the glycosylation of nicotine, UGT35B1 reduces its toxicity and promotes survival, highlighting a critical and previously unrecognized function of insect UGTs in xenobiotic metabolism.

### 3.4 Ugt35B1 confers protection against nicotine and flonicamid, but not imidacloprid or thiamethoxam

Having established that *Ugt35B1* can convert nicotine into nicotine *N*-glycoside, and with the recent development of several pyridine-based insecticides (Zakharychev and Martsynkevich, 2025), we compared three chemotypes: two neonicotinoids—imidacloprid, which contains a 6-chloro-3-pyridyl (i.e., 2-chloropyridine) ring, thiamethoxam, which lacks a pyridine ring— and flonicamid, a 4-(trifluoromethyl)nicotinamide (para-CF₃ on pyridine). To evaluate the substrate specificity of UGT35B1, we exposed CRISPR-knockout flies to these insecticides and measure either mortality or neurotoxic effects.

As expected, *Ugt35B1* KO flies were more sensitive to nicotine as previously shown (Fig. 4A). In addition, the *Ugt35B1* KO flies were more sensitive to flonicamid (Fig. 4B). In contrast, the *Ugt35B1* KO flies showed no difference in susceptibility to imidacloprid (Fig. 4C) or thiamethoxam (Fig. 4D) compared to controls. The simplest explanation of these results is that UGT35B1 requires a pyridine ring lacking substituents ortho of the N. Additional studies would be needed to more precisely determine the structural requirements on the pyridine ring (i.e. steric and electronic effects) for UGT35B1-mediated metabolism.

**Fig. 4.**
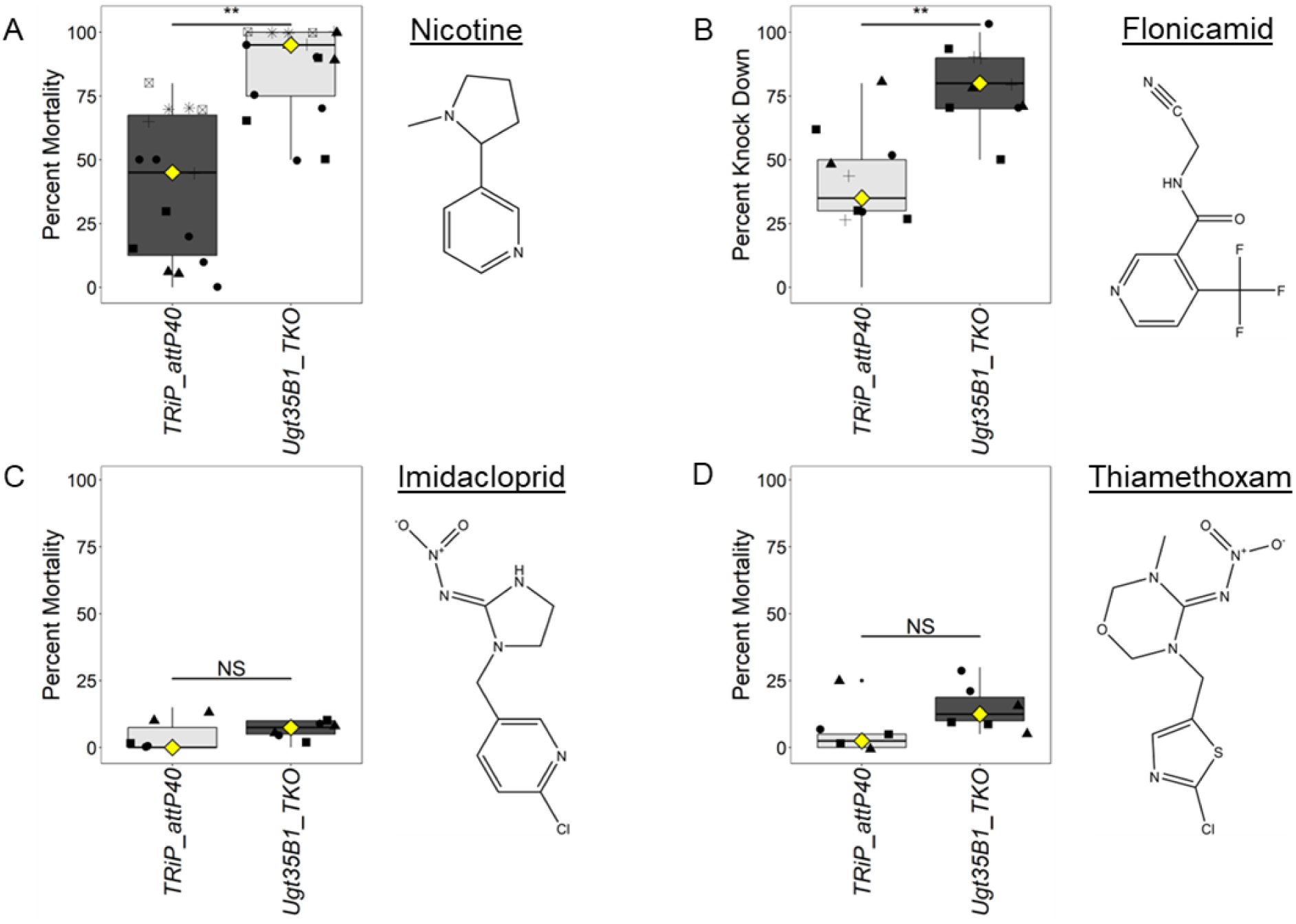
Toxicity of nicotine and flonicamid, but not imidacloprid nor thiamethoxam, is increased after *Ugt35B1* knockout. *Ugt35B1* TKO and its control were tested against other neonicotinoids. Box plots and chemical structures are shown for nicotine (A), flonicamid (B), imidacloprid (C), and thiamethoxam (D). Asterisks indicate significance between knockout and control lines. *p <0.05, **p < 0.01, ***p < 0.001.

## 4. Discussion

We found that adult *D. melanogaster* exposed to nicotine accumulated nicotine *N-*glycoside cotinine, and cotinine *N*-glycoside, implicating UDP-glycosyltransferases (UGTs) in nicotine metabolism. Likewise, the glycosylated form of cotinine is detected in cotinine-exposed flies. A targeted RNAi screen of most *D. melanogaster* UGTs identified a single gene, *Ugt35B1*, whose knockdown markedly increased susceptibility to nicotine. CRISPR-mediated knockout of *Ugt35B1* also increased toxicity of nicotine. LC-HRMS profiling of nicotine-exposed *Ugt35B1* knockouts confirmed that nicotine *N*-glycoside and cotinine *N*-glycoside were almost completely absent in *Ugt35B1* mutants. Together, these findings establish that *Ugt35B1* confers protection against nicotine. The *Ugt35B1* knockouts were also more sensitive to flonicamid, but not to imidacloprid nor thiamethoxam. As expected, there was glycosylation of flonicamid, but no glycosylation of imidacloprid detected by LCHRMS. These results are summarized graphically in supplemental figure S7.

A previous study identified a UGT null mutant (*Ugt35C1*) that enhanced the toxicity of nicotine to *D. melanogaster* larvae (Highfill et al., 2017; Macdonald and Highfill, 2020). In contrast, our screen did not detect any change in nicotine toxicity in adults following RNAi of *Ugt35C1*. This is not explained by life stage specific expression of the genes, as both are expressed in larvae and adults. It is possible that our RNAi for *Ugt35C1* was not effective at silencing expression, and we might have missed it for that reason. However, UGT35C1 does not appear to be a major cause of nicotine metabolism in adults, because our metabolism data suggest that UGT35B1 is responsible for the vast majority of nicotine conjugation in *D. melanogaster*.

Our finding of a single UGT that is responsible for production of most of the nicotine *N*-glycuronide is consistent with what has been found in other organisms. For example, in humans, the glucuronidation of nicotine and cotinine is primarily mediated by UGT2B10 (Chen et al., 2007; Kaivosaari et al., 2007), though other UGTs, including UGT1A1, UGT1A4, and UGT1A9, can catalyze these reactions in liver microsomes (Kuehl and Murphy, 2003; Nakajima et al., 2002). In *D. melanogaster*, glycosylation of nicotine and cotinine is strongly reduced, though not abolished, in *Ugt35B1* knockouts, suggesting that while UGT35B1 is the major enzyme involved, other UGTs may contribute minor activity.

While UGT35B1 was capable of metabolizing nicotine and flonicamid, no metabolism of imidacloprid was detected, likely do the presence of a Cl group ortho to the pyridine N. This suggests that UGT35B1 may not be particularly important for metabolism of neonicotinoids. For example, acetamiprid, nitenpyram, and thiacloprid, all both have a Cl group ortho to the pyridine N (like imidacloprid). Future quantitative structure activity relationship studies with UGT35B1 would be very valuable in discovering the chemical features that constrain UGT-mediated metabolism and will be informative as new pyridine insecticides come to market (Zakharychev and Martsynkevich, 2025).

Members of mammalian UGT1 and UGT2 families often act on multiple structurally diverse substrates (Meech et al., 2019). In arthropods, UGT substrate specificity has been less studied, though work in *Tetranychus urticae* shows that several UGTs can conjugate both plant metabolites and acaricides (Snoeck et al., 2019). Here, we show that UGT35B1 acts on nicotine (and cotinine) as well as flonicamid, but not on imidacloprid or thiamethoxam. Combined with its role in lipid homeostasis, this indicates that UGT35B1 is a multifunctional enzyme with the capacity to interact with both exogenous toxins and endogenous metabolites.

In conclusion, our findings identify UGT35B1 as the principal enzyme mediating nicotine glycosylation in *D. melanogaster.* Furthermore, UGT35B1 metabolizes cotinine (a metabolite of nicotine) and flonicamid. Collectively, our results show that nicotine metabolism in *D. melanogaster* is similar to what is found in mammals. Given the structural selectivity in UGT35B1 mediated metabolism we observed, further studies on UGT substrate specificity could inform both insect toxicology and the design of pest control agents less susceptible to metabolic detoxification.

## Acknowledgements

We thank Professor Gregory Loeb for valuable comments on this manuscript and Dr. Daniel Cordova for valuable discussions about the flonicamid bioassay. This research was supported by grants from USDA/NIFA (12898432 to JGS and NB) and NIH (R01AI148529 and R01AI148541 to NB) and NSF (IOS 2024252 to NB).

## Supplemental Figures

**Fig. S1.**
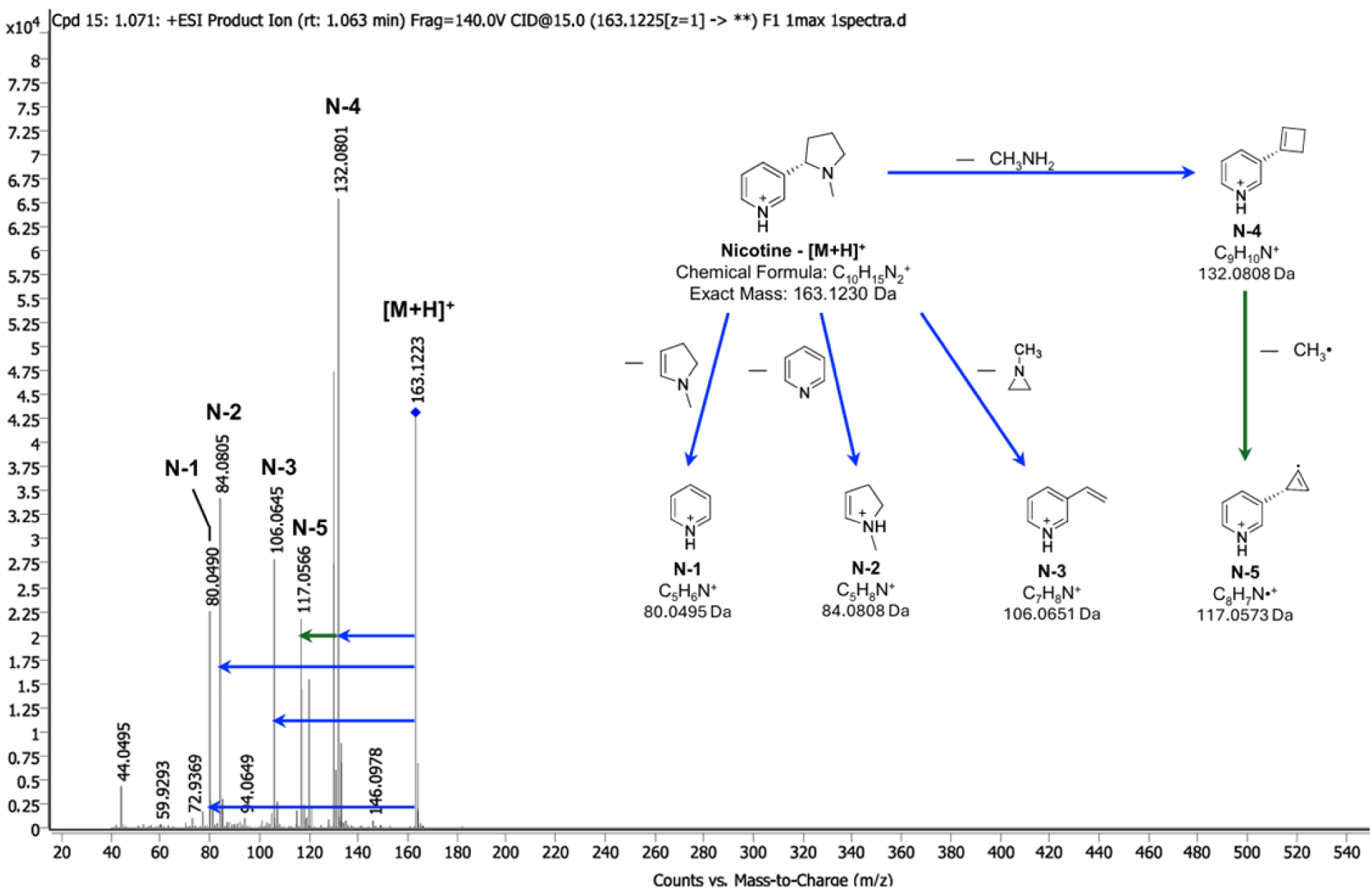
Tandem-MS product ion mass spectrum and putative fragmentation mechanism of [M+H]^+^ adduct of nicotine. MS^2^ (blue) and MS^3^ (green) fragments were identified based on the published putative nicotine fragmentation mechanism (Medana et al., 2016).

**Fig. S2.**
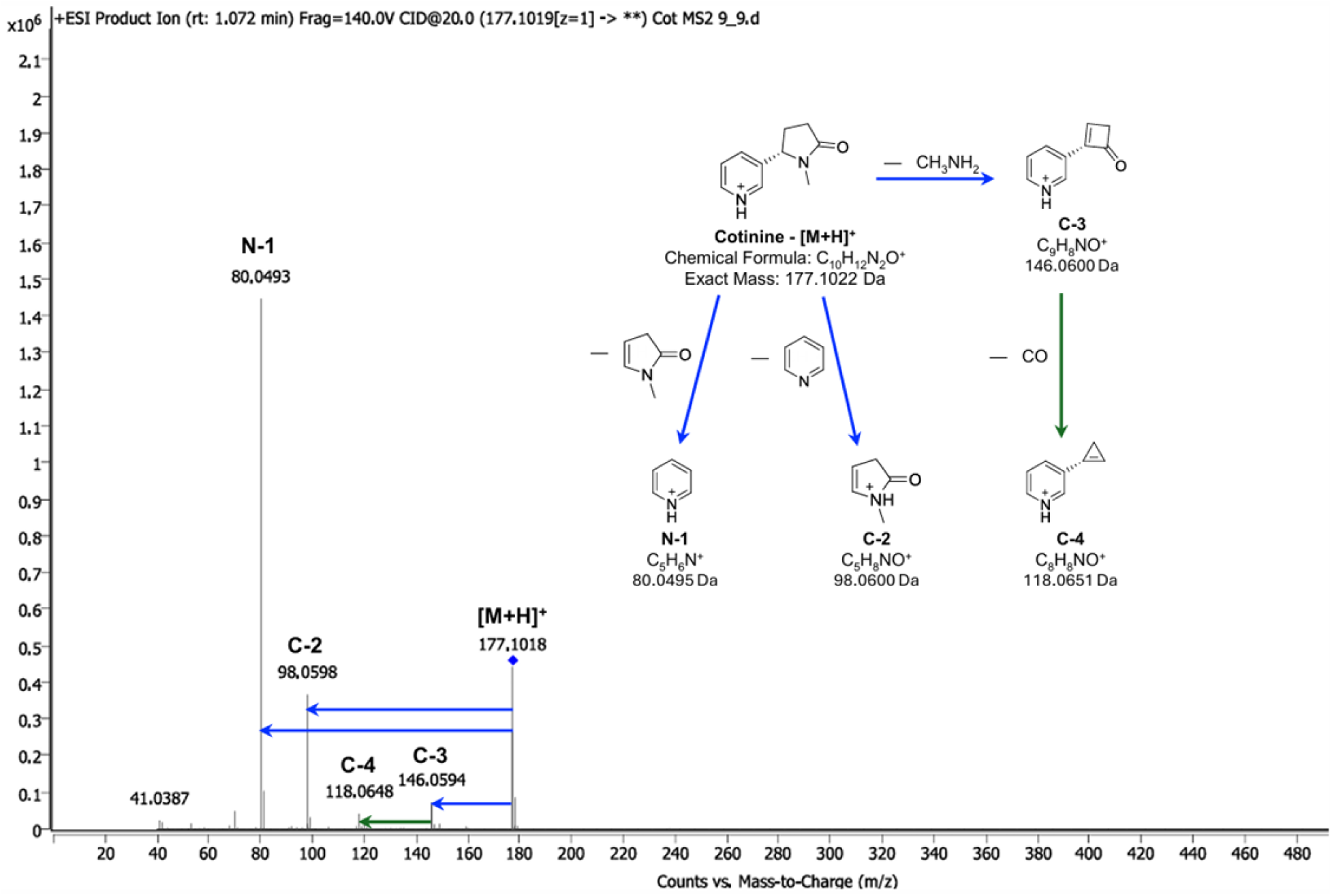
Tandem-MS product ion mass spectrum and putative fragmentation mechanism of [M+H]^+^ adduct of cotinine. MS^2^ (blue) and MS^3^ (green) fragments were identified based on the published putative cotinine fragmentation mechanism (Medana et al., 2016).

**Fig. S3.**
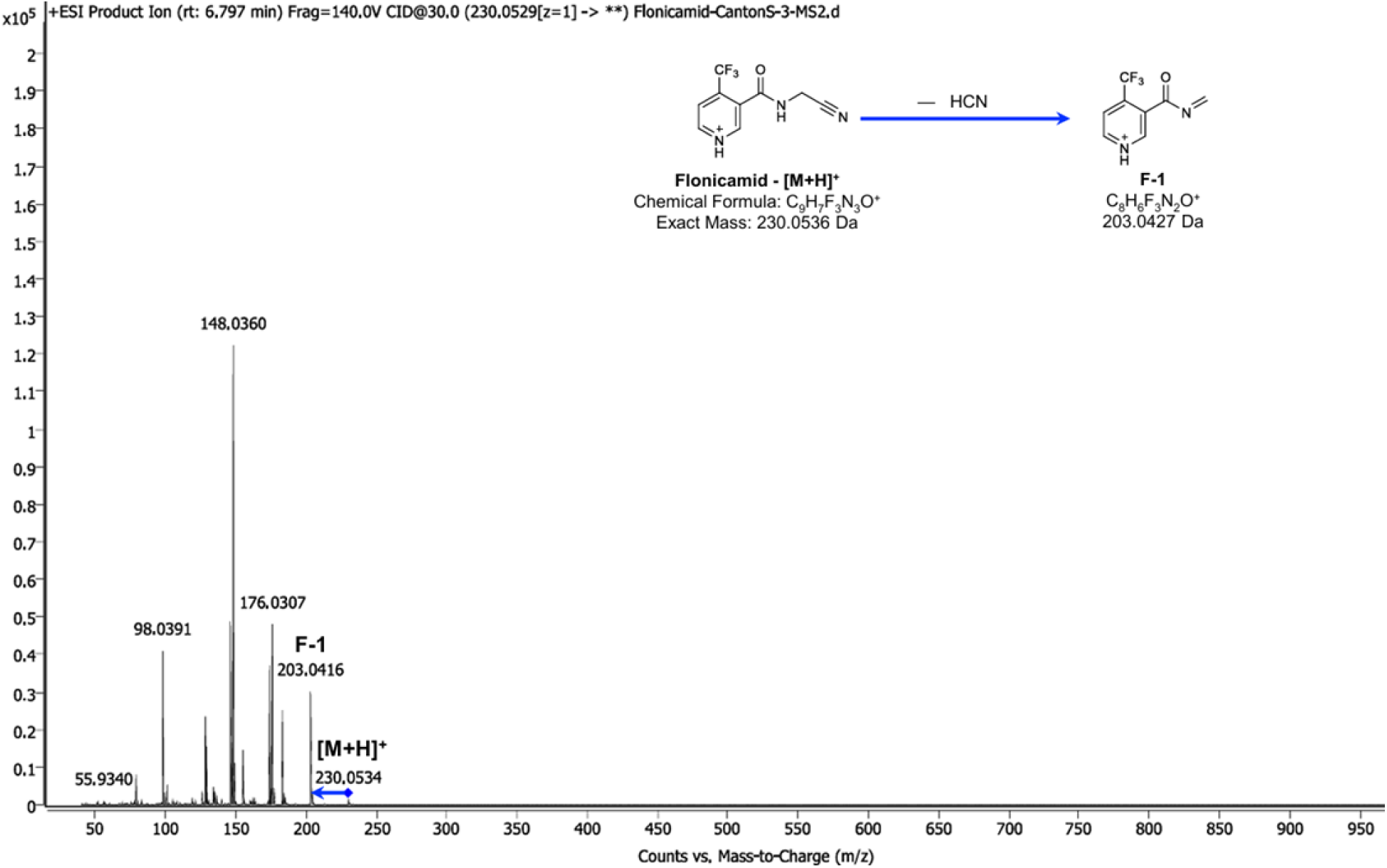
Tandem-MS product ion mass spectrum and putative fragmentation mechanism of [M+H]^+^ adduct of flonicamid.

**Fig. S4.**
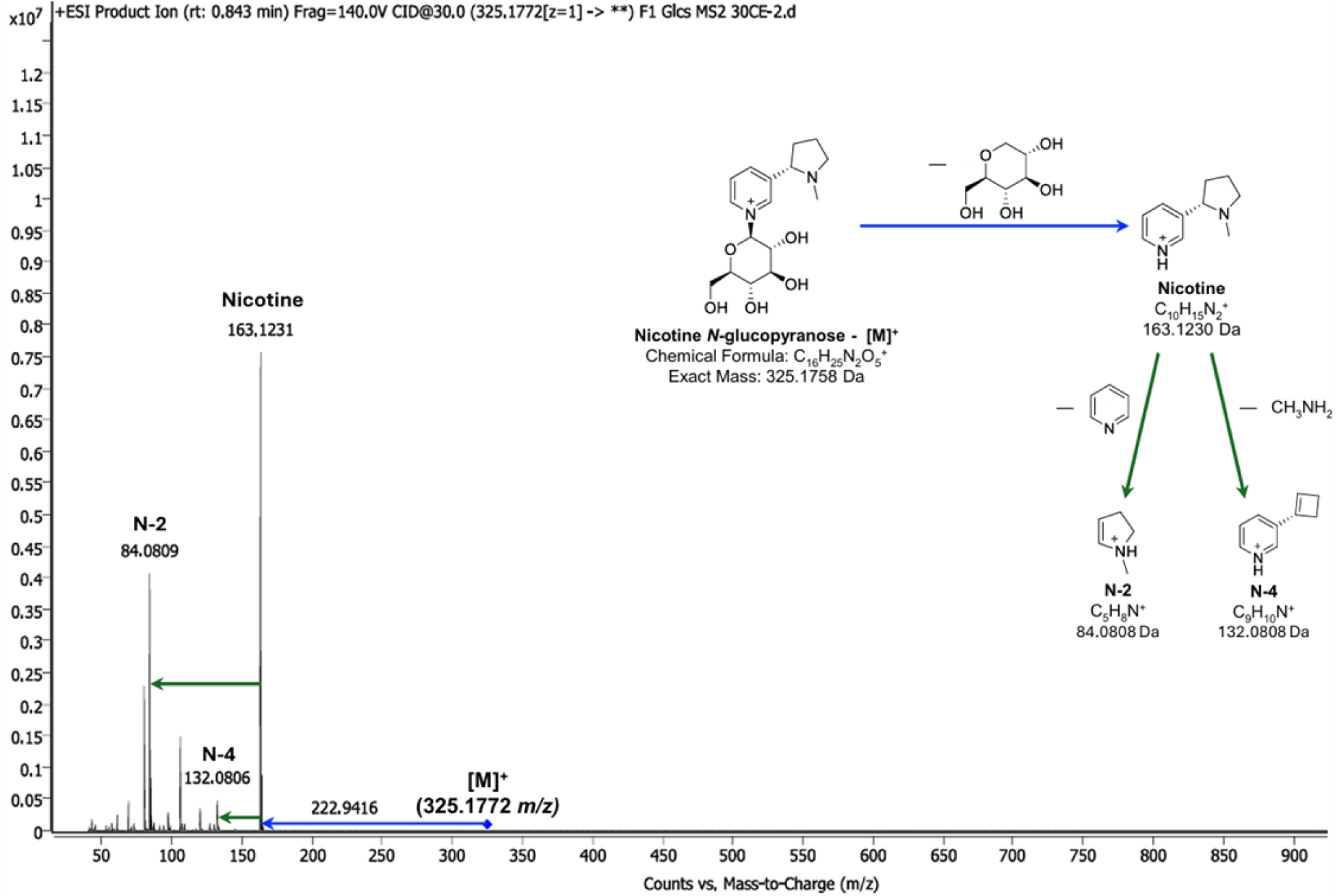
Tandem-MS product ion mass spectrum and putative fragmentation mechanism of [M]^+^ adduct of nicotine *N*-glycoside. Several fragments with the same exact mass were found in the tandem-MS profile of nicotine to support the proposed identification of nicotine *N*-glycoside.

**Fig. S5.**
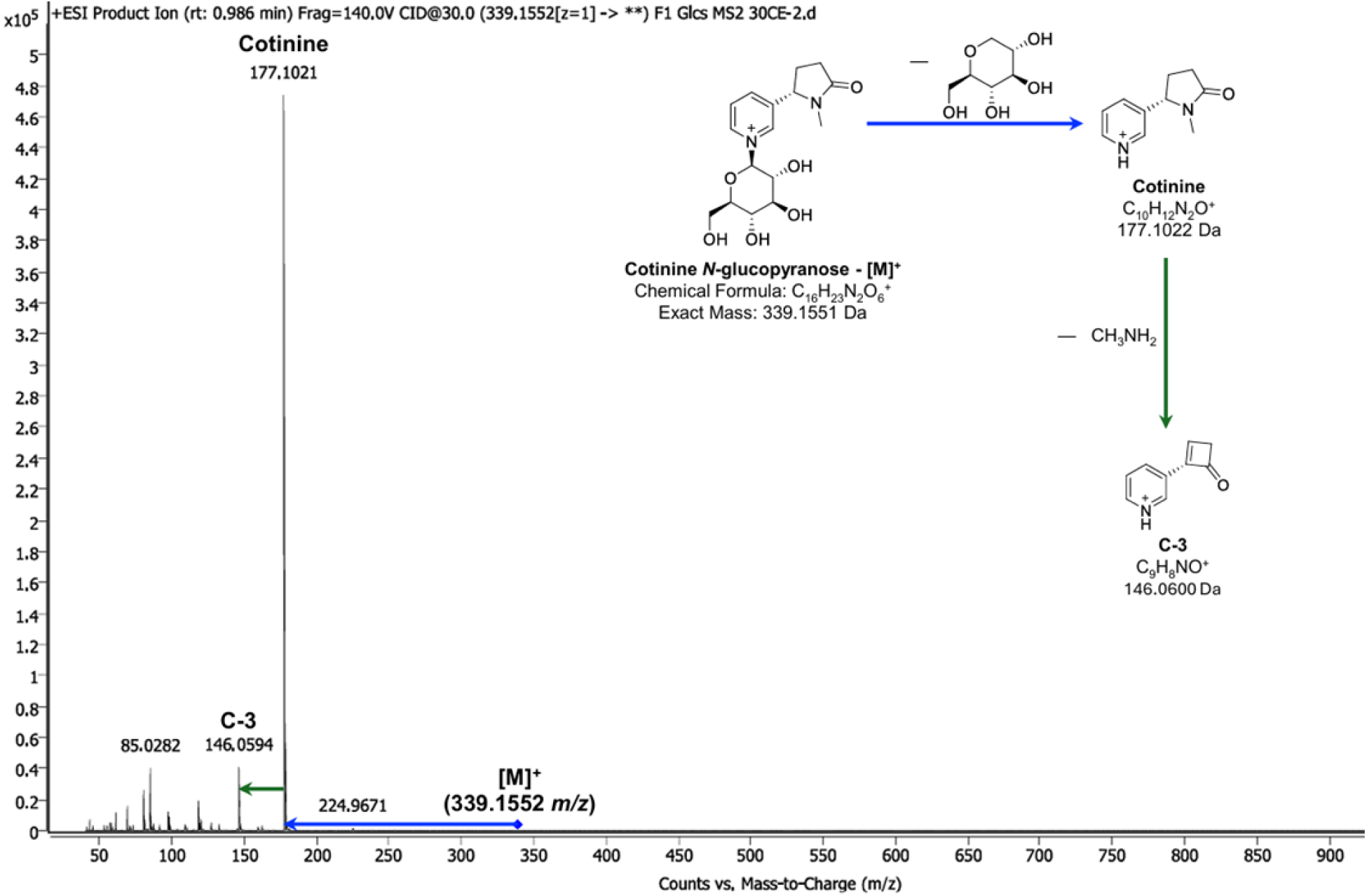
Tandem-MS product ion mass spectrum and putative fragmentation mechanism of [M]^+^ adduct of cotinine *N*-glycoside. Several fragments with the same exact mass were found in the tandem-MS profile of cotinine to support the proposed identification of cotinine *N*-glycoside.

**Fig. S6.**
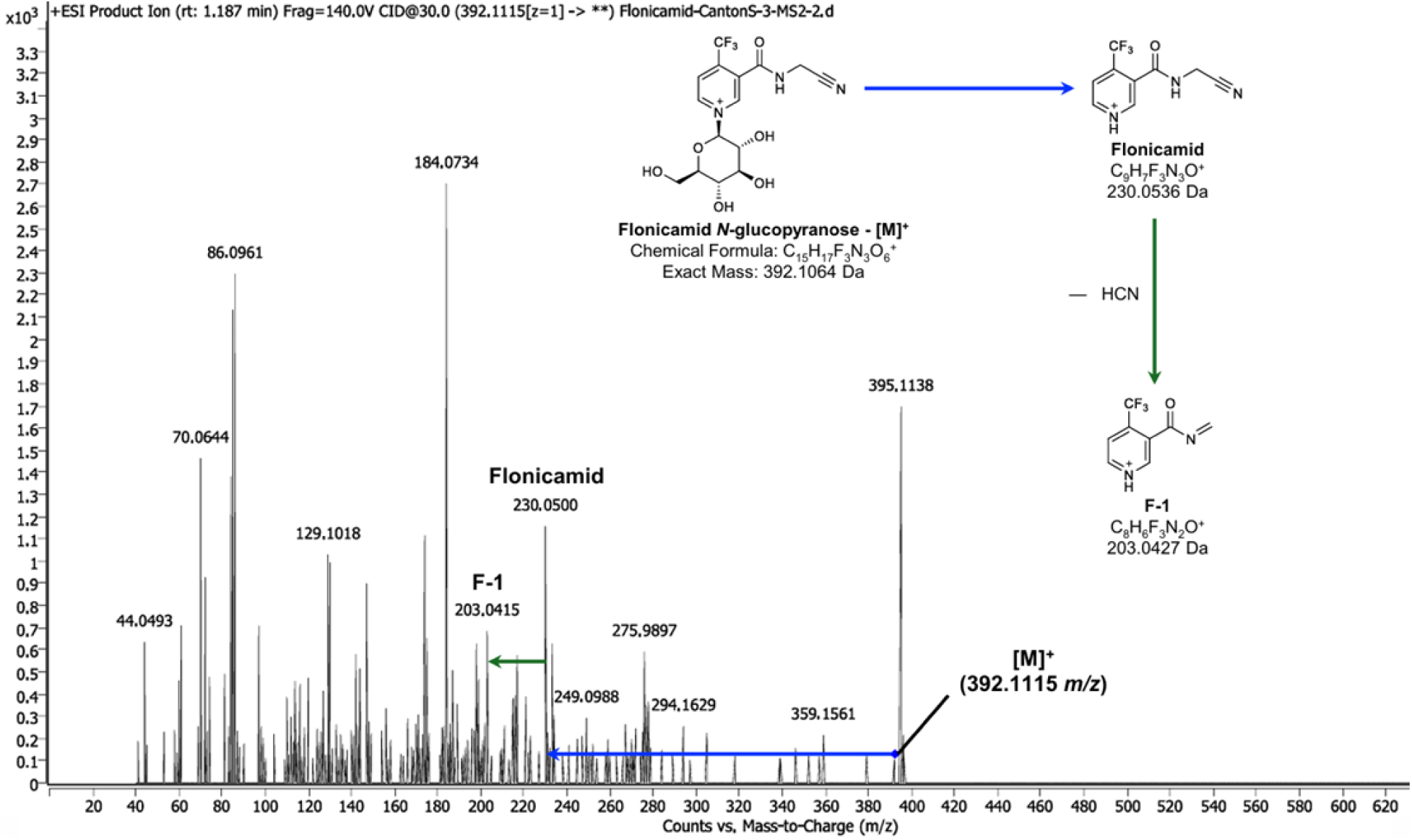
Tandem-MS product ion mass spectrum and putative fragmentation mechanism of [M]^+^ adduct of flonicamid *N*-glycoside. Several fragments with the same exact mass were found in the tandem-MS profile of flonicamid to support the proposed identification of flonicamid *N*-glycoside.

**Fig. S7.**
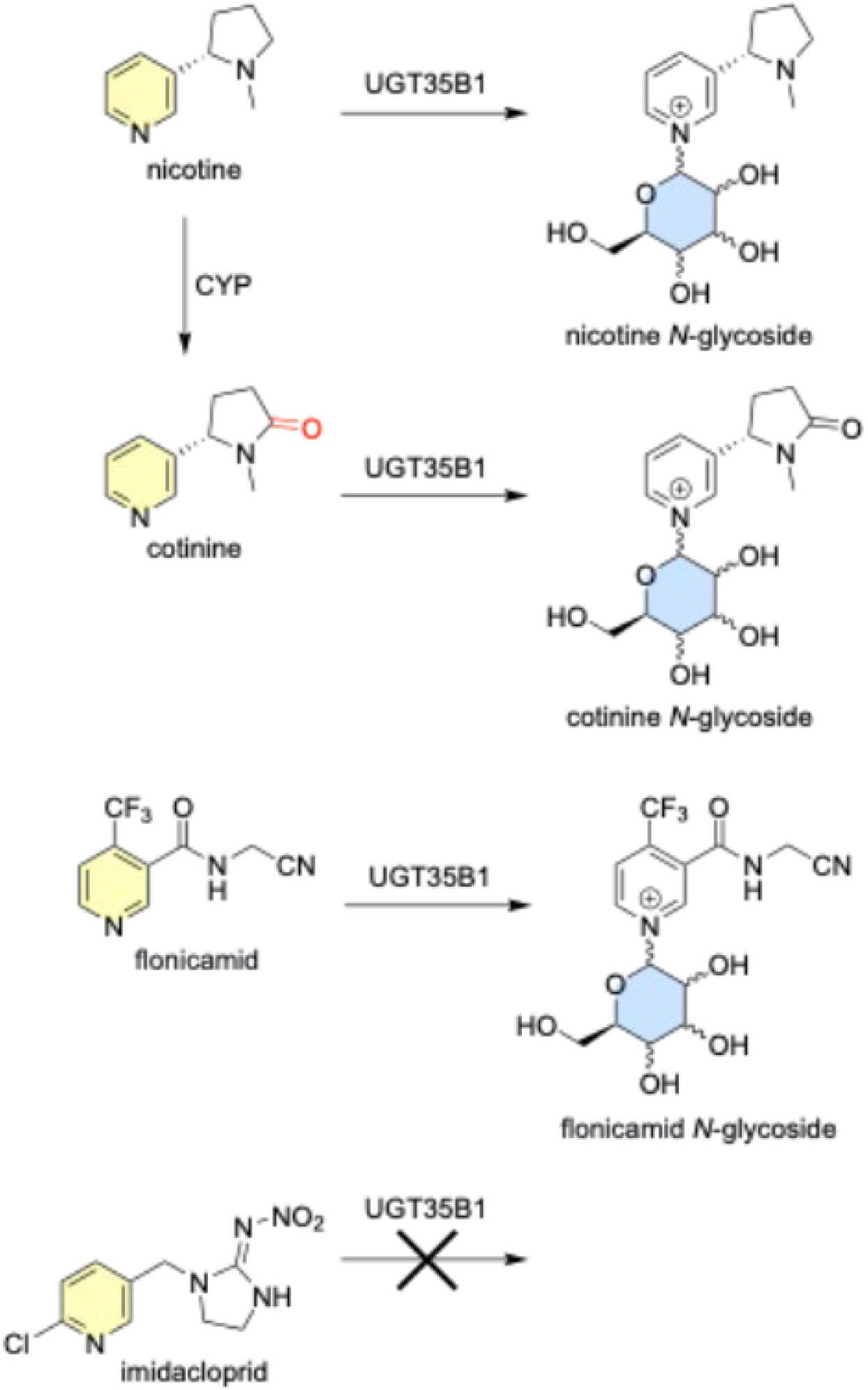
Imidacloprid is not glycosylated in *Drosophila*. Schematic showing the metabolic pathways by which nicotine would is transformed into cotinine, nicotine *N*-glycoside, and cotinine *N*-glycoside. Imidacloprid, as shown, is not metabolized into its glycosylated form in *Drosophila*.

**Table S1.** Drosophila strains used in this study.

| **Line Name** | **BDSC #** | **Line Type** |
| --- | --- | --- |
| UAS-Ugt35A1-IR | 67306 | TRiP RNAi |
| UAS-Ugt35B1-IR | 65083 | TRiP RNAi |
| UAS-Ugt35B1-IR | 64985 | TRiP RNAi |
| UAS-Ugt35D1-IR | 67305 | TRiP RNAi |
| UAS-Ugt35E1-IR | 62259 | TRiP RNAi |
| UAS-Ugt35E2-IR | 67307 | TRiP RNAi |
| UAS-Ugt36D1-IR | 62260 | TRiP RNAi |
| UAS-Ugt36E1-IR | 66346 | TRiP RNAi |
| UAS-Ugt37A1-IR | 62257 | TRiP RNAi |
| UAS-Ugt37A2-IR | 65082 | TRiP RNAi |
| UAS-Ugt37A3-IR | 64983 | TRiP RNAi |
| UAS-Ugt37C2-IR | 53931 | TRiP RNAi |
| UAS-Ugt37D1-IR | 64567 | TRiP RNAi |
| UAS-Ugt49B1-IR | 41967 | TRiP RNAi |
| UAS-Ugt49B2-IR | 64984 | TRiP RNAi |
| UAS-Ugt49C1-IR | 67302 | TRiP RNAi |
| UAS-Ugt50B3-IR | 51790 | TRiP RNAi |
| UAS-Ugt301D1-IR | 42657 | TRiP RNAi |
| UAS-Ugt302K1-IR | 64568 | TRiP RNAi |
| UAS-Ugt303A1-IR | 64982 | TRiP RNAi |
| UAS-Ugt304A1-IR | 62258 | TRiP RNAi |
| UAS-Ugt307A1-IR | 67301 | TRiP RNAi |
| UAS-Ugt316A1-IR | 80418 | TRiP RNAi |
| UAS-Ugt317A1-IR | 64981 | TRiP RNAi |
| TRiP attP2 | 36303 | TRiP RNAi |
| TRiP attP40 | 36304 | TRiP RNAi |
| Act5c-Gal4/cyo | 25374 | Driver |
| Act5c-Cas9 | 54590 | TRiP CRISPR |
| Ugt35B1_TKO | 83436 | TRiP CRISPR |

**Table S2.** HRMS data of the metabolites quantified in the samples.

| **Name** | **Retention time (min)** | **Precursor ion** | **Observed $m/z$** | **Calculated $m/z$** | **$\Delta m/z$ (ppm)** | **Formula** |
| --- | --- | --- | --- | --- | --- | --- |
| Nicotine | 1.06 | $[M+H]^+$ | 163.1223 | 163.1230 | -4.3 | $C_{10}H_{14}N_2$ |
| Cotinine | 1.12 | $[M+H]^+$ | 177.1020 | 177.1022 | -1.1 | $C_{10}H_{12}N_2O$ |
| Flonicamid | 6.79 | $[M+H]^+$ | 230.0533 | 230.0536 | -1.3 | $C_9H_6F_3N_3O$ |
| Nicotine *N*-glycoside | 0.84 | $[M]^+$ | 325.1772 | 325.1758 | 4.3 | $C_{16}H_{25}N_2O_5^+$ |
| Cotinine *N*-glycoside | 0.99 | $[M]^+$ | 339.1552 | 339.1551 | 0.3 | $C_{16}H_{23}N_2O_6^+$ |
| Flonicamid *N*-glycoside | 1.17 | $[M]^+$ | 392.1044 | 392.1064 | -5.1 | $C_{15}H_{17}F_3N_3O_6^+$ |

## References

Ahmad, S., Forgash, A.J., 1976. Nonoxidative enzymes in the metabolism of insecticides. Drug Metab. Rev. 5, 141–164.

Ahn, S.-J., Badenes-Perez, F.R., Reichelt, M., Svatos, A., Schneider, B., Gershenzon Jonathan, Heckel, D.G., 2011. Metabolic detoxification of capsaicin by UDP-glycosyltransferase in three *Helivoerpa* species. Arch. Insect Biochem. Physiol. 78, 104–118.

Ahn, S.-J., Vogel, H., Heckel, D.G., 2012. Comparative analysis of the UDP-glycosyltransferase multigene family in insects. Insect Biochem. Mol. Biol. 42, 133–147.

Chen, G., Blevins-Primeau, A.S., Dellinger, R.W., Muscat, J.E., Lazarus, P., 2007. Glucuronidation of nicotine and cotinine by UGT2B10: loss of function by the UGT2B10 codon 67 (Asp>Tyr) polymorphism. Cancer Res. 67, 9024–9029.

du Rand, E., Pirk, C.W.W., Nicolson, S.W., Apostolides, Z., 2017. The metabolic fate of nectar nicotine in worker honey bees. J. Insect Physiol. 98, 14–22.

Highfill, C.A., Tran, J.H., Nguyen, S.K.T., Moldenhauer, T.R., Wang, X., Macdonald, S.J., 2017. Naturaly segregating variation at *Ugt86Dd* contributes to nicotine resistance in *Drosophila melanogaster*. Genetics 207, 311–325.

Jeschke, P., Nauen, R., 2008. Neonicotinoids - from zero to hero in insecticide chemistry. Pest Manag. Sci. 64, 1084–1098.

Kaivosaari, S., Toivonen, P., Hesse, L.M., Koskinen, M., Court, M.H., Finel, M., 2007. Nicotine glucuronidation and the human UDP-glucuronosyltransferase UGT2B10. Mol. Pharmacol. 72, 761–768.

Kinareikina, A.G., Silivanova, E.A., 2024. The role of UDP-glycosyltransferases in xenobiotic metabolism. J. Evol. Biochem. Physiol. 60, 1920–1942.

Krempl, C., Sporer, T., Reichelt, M., Ahn, S.-J., Heidel-Fischer, H., Vogel, H., Heckel, D.G., Joußen, N., 2016. Potential detoxification of glossypol by UDP-glycosyltransferases in the two Heliothine moth species *Helicoverpa armigera* and *Heliothis virescens*. Insect Biochem. Mol. Biol. 71, 49–57.

Kuehl, G.E., Murphy, S.E., 2003. *N*-glucuronidation of nicotine and cotinine by human liver micorsomes and heterologously expressed UDP-glucuronosyltransferases. Drug Metab. Dispos. 31, 1361–1368.

Macdonald, S.J., Highfill, C.A., 2020. A naturally-occurring 22-bp coding deletion in *Ugt86Dd* reduces nicotine resistance in *Drosophila melanogaster*. BMC Res. Notes 13.

McIndoo, N.E., 1916. Effects of nicotine as an insecticide. J. Agric. Res. 7, 89–122.

McKinlay, R.G., Spaull, A.M., Straub, R.W., 1992. Pests of solanaceous crops, in: Vegetable Crop Pests. Palgrave Macmillan, London, pp. 263–326.

Medana, C., Santoro, V., Bello, F.D., Sala, C., Pazzi, M., Sarro, M., Calza, P., 2016. Mass spectrometric fragmentation and photocatalytic transformation of nicotine and cotinine. Rapid Comm Mass Spectrom 30, 2617–2627.

Meech, R., Hu, D.G., McKinnon, R.A., Mubarokah, S.N., Haines, A.Z., Nair, P.C., Rowland, A., Mackenzie, P.I., 2019. The UDP-glycosyltransferase (UGT) superfamily: new members, new functions, and novel paradigms. Physiol Rev 99, 1153–1222. 10.1152/physrev.00058.2017

Morris, C.E., 1983. Uptake and metabolism of nicotine by the CNS of a nicotine-resistant insect, the tobacco hornworm (*Manduca sexta*). J. Insect Physiol. 29, 807–817.

Murphy, S.E., 2021. Biochemistry of nicotine metabolism and its relevance to lung cancer. J. Biol. Chem. 296.

Nakajima, M., Tanaka, E., Kwon, J.-T., Yokoi, T., 2002. Characterization of nicotine and cotinine *N*-glucuronidations in human liver micosomes. Drug Metab. Dispos. 30, 1484–1490.

Qiao, X., Zhang, Xiaoyu, Zhou, Z., Guo, L., Wu, W., Ma, S., Zhang, Xinzhong, Montell, C., Huang, J., 2022. An insecticide target in mechanoreceptor neurons. Sci. Adv. 23. 10.1126/sciadv.abq3132

Robert, C.A.M., Zhang, X., Machado, R.A.R., Schirmer, S., Lori, M., Mateo, P., Erb, M., Gershenzon, J., 2017. Sequestration and activation of plant toxins protect the western corn rootworm from enemies at multiple trophic levels. eLife 6.

Saremba, B.M., Murch, S.J., Tymm, F.J.M., Rheault, M.R., 2018. The metabolic fate of dietary nicotine in the cabbage looper, *Trichoplusia ni* (Hubner). J. Insect Physiol. 109, 1–10.

Scott, J.G., 2025. Insect UDP-glycosyltransferases and xenobiotic metabolism. Pestic Biochem Physiol.

Self, L.S., Guthrie, F.E., Hodgson, E., 1964. Metabolism of nicotine by tobacco-feeding insects. Nature 204, 300–301.

Snoeck, S., Pavlidi, N., Pipini, D., Vontas, J., Dermauw, W., Leeuwen, T.V., 2019. Substrate specificity and promiscuity of horizontally transferred UDP-glycosyltransferases in the generalit herbivore *Tetranychus urticae*. Insect Biochem. Mol. Biol. 109, 116–127.

Snyder, M.J., Walding, J.K., Feyereisen, R., 1994. Metabolic fate of the allelochemical nicotine in tobacco hornworm *Manduca sexta*. Insect Biochem. Mol. Biol. 24, 837–846.

Wouters, F.C., Reichelt, M., Glauser, G., Bauer, E., Erb, M., Gershenzon, J., Vassao, D.G., 2014. Reglucosylation of the benzoxazinoid DIMBOA with inversion of the stereochemical configuration is a detoxification strategy in Lepidopteran herbivores. Angew. Chem. Int. Ed. 53, 11320–11324.

Yamamoto, I., 1999. Nicotine to nicotinoids: 1962-1997, in: Nicotinoid Insecticides and the Nicotinic Acetylcholine Receptor. Springer, Tokyo, pp. 3–27.

Yildiz, D., 2004. Nicotine, its metabolism and an overview of its biological effects. Toxicon 43, 619–632.

Zakharychev, V.V., Martsynkevich, A.M., 2025. Development of novel pyridine-based agrochemicals: A review. Adv. Agrochem 4, 30–48.

Ziemke, T., Wang, P., Duplais, C., 2024. The fate of a *Solanum* steroidal alkaloid toxin in the cabbage looper (*Trichoplusia ni*). Insect Biochem. Mol. Biol. 175, 104205.

